## Supplementary material for "H-NS is a bacterial transposon capture protein": Table S1

Table S1: Changes in gene expression in grey colony derivatives compared to wild types cells

**Locus	Name	Function	Log fold change	P value**

*Up regulated genes*

ABUW_1703	hypothetical protein	2.41	9.21E-03

ABUW_2112	DUF2726 domain protein	2.41	2.29E-02

ABUW_1766	IS*Aba*13	insertion sequence	2.14	1.14E-09

ABUW_3803	IS*Aba*13	insertion sequence	2.07	3.78E-09

*Down regulated genes*

ABUW_2006	major capsid protein	-2.12	5.09E-06

ABUW_2017	major capsid protein	-2.25	1.87E-04

ABUW_2028	major capsid protein	-2.54	2.80E-07

ABUW_1769	PEGA domain protein	-2.80	2.37E-06

ABUW_3823	*weeI*	sugar transferase	-2.94	1.82E-11

ABUW_1641	hypothetical protein	-3.25	1.35E-05

ABUW_3822	*weeH*	acetyl transferase	-3.43	2.08E-13

ABUW_0304	*pilA*	pilin	-5.49	1.22E-31
