## Supplementary material for "H-NS is a bacterial transposon capture protein": Table S3

**Table S3: Mobile genetic elements containing transposition hotspots dependent on H-NS**

**Mobile element	Chromosomal position	Genes with H-NS dependent**

**transposition hotspots**

*Chromosomal elements*

Aeromonas phage SW69-9	318756-329326	0

Escherichia phage vB_EcoM_ECO1230-10	545886-581029	1

Acinetobacter phage Bphi-B1251	736407-769903	3

Acinetobacter phage Bphi-B1251	776101-793726	0

Acinetobacter phage Bphi-B1251	1289765-1330967	6

Acinetobacter phage Bphi-B1251	1314667-1338244	2

Acinetobacter phage Ab105-2phi	1391199-1409865	0

Acinetobacter phage Ab105-2phi	1393236-1416423	0

Enterobacteria phage Ike	1985246-2029746	1

Acinetobacter phage Ab105-1phi	2650307-2669540	7

Moraxella phage Mcat16	3031838-3049068	1

*Extra chromosomal elements*

83.61 kb plasmid	N.A.	18

8.73 kb plasmid	N.A.	2

1.97 kb plasmid	N.A.	0
