## Supplementary material for "H-NS is a bacterial transposon capture protein": Table S4

**Table S4:** Strains and plasmids

**Name	Description	Source**

***A. baumannii* strains**

AB5075	Highly virulent and drug resistant isolate from an osteomyelitis	(1)

tibial infection.

AB5075::*gtr52*::IS*Aba*13	Naturally occurring “grey” AB5075 derivative with an insertion sequence	This work

disrupting the *gtr52* gene.

AB5075::*ompW*::IS*Aba*13	Derivative of AB5075 with an insertion sequence disrupting the	This work

*ompW* gene, generated by scarless genome editing.

AB5075 *hns*::T26	Derivative of AB5075 with a T26 transposon in *hns*. Tet^R^.	(2)

***E. coli* strains**

JCB387	Used for general plasmid DNA manipulation. Δ*nirB*Δ*lac*	.	(3)

DH5α	Used for general plasmid DNA manipulation. *fhuA*2Δ(*argF*-*lacZ*)U169	NEB	*phoA* *glnV*44 Φ80Δ(*lacZ*)M15 *gyrA*96 *recA*1 *relA*1 *endA*1 *thi*-1 *hsdR*17.

T7 Express	Used to overexpress *A. baumannii* H-NS from pJ414.	NEB

MG1655	Used for native Tn-seq analysis of *insH3*.	(4)

**Plasmids**

pVRL1Z	High copy number plasmid, contains *parE*2-*paaA*2 toxin-antitoxin	(5)

system. Zeo^R^. Used to express H-NS-39 in *A. baumannii*.

pVLR2Z	High copy number plasmid, contains *parE*2-*paaA*2 toxin-antitoxin	(5)

system. Zeo^R^. Has an arabinose inducible promoter.

pMHL-2	Template for PCR used to make DNA fragments for genome editing.	(6)

Contains *apra*R::*sacB* counter selection and resistance cassette.

pJ414	Used to overexpress *A. baumannii* H-NS in *E. coli*.	ATUM
