## Supplementary material for "H-NS is a bacterial transposon capture protein": Table S5

**Table S5:** Oligonucleotides

**Name	Description	Sequence (5' to 3')**

**for generation of *ompW*::IS*Aba*13**

P5	To amplify *apraR*::*sacB* cassette	cgactcactatagggcgaattgggccgctttccagtcgggaaacctg

P6	To amplify *apraR*::*sacB* cassette	catatgccaccgacccgagcaaaccccgccagggttttcccagtcacgac

P62	To create fragment 1	cagcagtcacataatagatagc

P63	To create fragment 1	ggcccaattcgccctatagtgagtcgattggcaagtaaaatttggg

P64	To create fragment 2	gggtttgctcgggtcggtggcatatgggctttgttgcacaaagatttaaaag

P65	To create fragment 2	attggcaagggctttgttgcacaaacctatctc

P66	To create fragment 3	caacaaagcccttgccaattaccagca

P71	To create fragment 3	cgttatgcgcaatgtccagt

P70	To amplify Gibson assembly product	taagccatcaagcaaagtgag

P74	To amplify Gibson assembly product ctcagagctaataagtgactg

and create fragment used in second

recombination

P72	For creating fragment used in second	ccgtactaccttctacacggt

recombination

P157	For creating fragment used in second acttgccaatggctttgttgcacaaagatttaaaagttaag

recombination

P158	For creating fragment used in second caacaaagccattggcaagtaaaatttggg

recombination

**for generation of *hns-*3x-FLAG**

prSL1	To create fragments 1 and 5	ggctattaattgctgagcaagctttg

prSL11	To create fragment 2	ggtgcaaaacttgaagatttcttaatcgctactgactacaaagaccatgac

prSL12	To create fragment 2	gactgggaaaaccctggcgttacttgtcatcgtcatccttgtaatcg

prSL13	To create fragment 1	caccgtcatggtctttgtagtcagtagcgattaagaaatcttcaagttttgcacc

prSL14	To create fragment 3	cgattacaaggatgacgatgacaagtaacgccagggttttcccagtc

prSL23	To create fragment 6	gattacaaggatgacgatgacaagtaattgtattgcctcttaaaaagccaagcg

prSL24	To create fragment 5	cgcttggctttttaagaggcaatacaattacttgtcatcgtcatccttgtaatc

prSL3	To create fragment 4	ggtttcccgactggaaagcgttgtattgcctcttaaaaagccaagcgattc

prSL4	To create frarments 4 and 5	gtggacgttgatgattcaataaagcc

prSL6	To create fragment 3	cgcttggctttttaagaggcaatacaacgctttccagtcgggaaacc

prSL9	To amplify Gibson assembly product	gcaactagccaacaactcaaaaacc

prSL10	To amplify Gibson assembly product	gttgtatggtcatcacttgatcaccac

**for cloning *A. baumannii hns* in pJ414**

P129	To amplify codon optimised AB5075 gcaagccatatgaaaccggacattagc

*hns* for cloning in pJ414. *Nde*I

restriction site underlined.

P146	To amplify codon optimised AB5075 gcaggtctcgagttaaatcaggaaatc

*hns* for cloning in pJ414. *Xho*I

restriction site underlined.

**for constructing *hns*-39 and cloning in pVRL1Z**

P134	To create fragment 1, *Xho*I site	ccccctcgagataaatattaagaaaatatattacaattataattactaatg

underlined

P135	To create fragment 1	tgatcttttttcattaataaatactccagtcttac

P136	To create fragment 2	tttattaatgaaaaaagatcaagcaatcg

P137	To create fragment 2, *Pst*I underlined cgggctgcagttatgttgttttcttacgtttttg

**for *in vitro* DNA bridging assays**

P162	To create IS*Aba*13 bait fragment	biotin-cttattaaatggctttgttgcac

P163	To create IS*Aba*13 bait fragment	taatttaataaggctttgttgcac

P171	To create T6SS prey fragment	caacacaactttcattcc

P172	To create T6SS prey fragment	agggtatctatatcagcca

**for native Tn-seq with *A. baumannii***

Adapter 1.2	Anneals with adapter 2.2	taccacgacca-NH_2_

Adapter 2.2	Anneals with adapter 1.2	atgatggccggtggatttgtgtggtcgtggtat

JelAP1	For 1st PCR, binds adapter 2.2	atgatggccggtggatttgtg

sequence

IS*Aba*13_out	For 1st PCR, binds near to the 5'end	caaagccaagtcaatgagattcatgc

of IS*Aba*13

Staggered_1	For 2nd hemi-nested PCR of P5 end,	aatgatacggcgaccaccgagatctacactctttccctacacgacgctcttccgatc*t*g

binding site to IS*Aba*13 underlined.	accacatacccgagttgtcac

T in italic added for heterogeneity.

Staggered_2	For 2nd hemi-nested PCR of P5 end,	aatgatacggcgaccaccgagatctacactctttccctacacgacgctcttccgatc*tt*

binding site to IS*Aba*13 underlined.	gaccacatacccgagttgtcac

Ts in italic added for heterogeneity.

Staggered_3	For 2nd hemi-nested PCR of P5 end,	aatgatacggcgaccaccgagatctacactctttccctacacgacgctcttccgatc

binding site to IS*Aba*13 underlined.	*tgata*gaccacatacccgagttgtcac

TGATA in italic added for

heterogeneity.

Staggered_4	For 2nd hemi-nested PCR of P5 end,	aatgatacggcgaccaccgagatctacactctttccctacacgacgctcttccgatc

binding site to IS*Aba*13 underlined.	*tatcta*gaccacatacccgagttgtcac

TATCTA in italic added for

heterogeneity.

AP1_P7_tagged_1	For 2nd hemi-nested PCR of P7 end,	caagcagaagacggcatacgagat*atcacg*gtgactggagttcagacgtgtgctctt

binding site to adaptor underlined.	ccgatctgtcaatgatggccggtggatttgtg

Barcode: CGTGAT in italic.

AP1_P7_tagged_2 For 2nd hemi-nested PCR of P7 end, caagcagaagacggcatacgagat*cgatgt*gtgactggagttcagacgtgtgctctt

binding site to adaptor underlined.	ccgatctgtcaatgatggccggtggatttgtg

Barcode: ACATCG in italic.

AP1_P7_tagged_3 For 2nd hemi-nested PCR of P7 end, caagcagaagacggcatacgagat*ttaggc*gtgactggagttcagacgtgtgctctt

binding site to adaptor underlined.	ccgatctgtcaatgatggccggtggatttgtg

Barcode: GCCTAA in italic.

AP1_P7_tagged_4 For 2nd hemi-nested PCR of P7 end, caagcagaagacggcatacgagat*tgacca*gtgactggagttcagacgtgtgctctt

binding site to adaptor underlined.	ccgatctgtcaatgatggccggtggatttgtg

Barcode: TGGTCA in italic.

AP1_P7_tagged_5 For 2nd hemi-nested PCR of P7 end, caagcagaagacggcatacgagat*acagtg*gtgactggagttcagacgtgtgctctt

binding site to adaptor underlined.	ccgatctgtcaatgatggccggtggatttgtg

Barcode: CACTGT in italic.

AP1_P7_tagged_6 For 2nd hemi-nested PCR of P7 end, caagcagaagacggcatacgagat*gccaat*gtgactggagttcagacgtgtgctctt

binding site to adaptor underlined.	ccgatctgtcaatgatggccggtggatttgtg

Barcode: ATTGGC in italic.

AP1_P7_tagged_7 For 2nd hemi-nested PCR of P7 end, caagcagaagacggcatacgagat*gctatg*gtgactggagttcagacgtgtgctctt

binding site to adaptor underlined.	ccgatctgtcaatgatggccggtggatttgtg

Barcode: CATAGC in italic.

AP1_P7_tagged_8 For 2nd hemi-nested PCR of P7 end, caagcagaagacggcatacgagat*agctag*gtgactggagttcagacgtgtgctctt

binding site to adaptor underlined.	ccgatctgtcaatgatggccggtggatttgtg

Barcode: CTAGCT in italic.

AP1_P7_tagged_9 For 2nd hemi-nested PCR of P7 end, caagcagaagacggcatacgagat*gtcgaa*gtgactggagttcagacgtgtgctctt

binding site to adaptor underlined.	ccgatctgtcaatgatggccggtggatttgtg

Barcode: TTCGAC in italic.

AP1_P7_tagged_10 For 2nd hemi-nested PCR of P7 end,caagcagaagacggcatacgagat*tacgag*gtgactggagttcagacgtgtgctctt

binding site to adaptor underlined.	ccgatctgtcaatgatggccggtggatttgtg

Barcode: CTCGTA in italic.

**for native Tn-seq with *E. coli***

*insH3*_out	For 1st PCR, binds near to the 5'end	gataacgccttaaatggcgaagaaac

of *insH3*

Staggered_1	For 2nd hemi-nested PCR of P5 end,	aatgatacggcgaccaccgagatctacactctttccctacacgacgctcttccgatc*t*g

binding site to *insH3* underlined.	ggagaaaaaatcggctcaaacatg

T in italic added for heterogeneity.

Staggered_2	For 2nd hemi-nested PCR of P5 end,	aatgatacggcgaccaccgagatctacactctttccctacacgacgctcttccgatc*tt*

binding site to *insH3* underlined.	gggagaaaaaatcggctcaaacatg

Ts in italic added for heterogeneity.

Staggered_3	For 2nd hemi-nested PCR of P5 end,	aatgatacggcgaccaccgagatctacactctttccctacacgacgctcttccgatc

binding site to *insH3* underlined.	*tgata*gggagaaaaaatcggctcaaacatg

TGATA in italic added for

heterogeneity.

Staggered_4	For 2nd hemi-nested PCR of P5 end,	aatgatacggcgaccaccgagatctacactctttccctacacgacgctcttccgatc

binding site to *insH3* underlined.	*tatcta*gggagaaaaaatcggctcaaacatg

TATCTA in italic added for

heterogeneity.
